# Exomap1 mouse: a transgenic model for *in vivo* studies of exosome biology

**DOI:** 10.1101/2023.05.29.542707

**Authors:** Francis K. Fordjour, Sarah Abuelreich, Xiaoman Hong, Emeli Chatterjee, Valeria Lallai, Martin Ng, Andras Saftics, Fengyan Deng, Natacha Carnel-Amar, Hiroaki Wakimoto, Kazuhide Shimizu, Malia Bautista, Tuan Anh Phu, Ngan K. Vu, Paige C. Geiger, Robert L. Raffai, Christie D. Fowler, Saumya Das, Lane K. Christenson, Tijana Jovanovic-Talisman, Stephen J. Gould

## Abstract

Exosomes are small extracellular vesicles (sEVs) of ∼30-150 nm in diameter that have the same topology as the cell, are enriched in selected exosome cargo proteins, and play important roles in health and disease. To address large unanswered questions regarding exosome biology *in vivo*, we created the *exomap1* transgenic mouse model. In response to Cre recombinase, *exomap1* mice express HsCD81mNG, a fusion protein between human CD81, the most highly enriched exosome protein yet described, and the bright green fluorescent protein mNeonGreen. As expected, cell type-specific expression of Cre induced the cell type-specific expression of HsCD81mNG in diverse cell types, correctly localized HsCD81mNG to the plasma membrane, and selectively loaded HsCD81mNG into secreted vesicles that have the size (∼80 nm), topology (outside out), and content (presence of mouse exosome markers) of exosomes. Furthermore, mouse cells expressing HsCD81mNG released HsCD81mNG-marked exosomes into blood and other biofluids. Using high-resolution, single-exosome analysis by quantitative single molecule localization microscopy, we show here that that hepatocytes contribute ∼15% of the blood exosome population whereas neurons contribute <1% of blood exosomes. These estimates of cell type-specific contributions to blood EV population are consistent with the porosity of liver sinusoidal endothelial cells to particles of ∼50-300 nm in diameter, as well as with the impermeability of blood-brain and blood-neuron barriers to particles >5 nm in size. Taken together, these results establish the *exomap1* mouse as a useful tool for *in vivo* studies of exosome biology, and for mapping cell type-specific contributions to biofluid exosome populations. In addition, our data confirm that CD81 is a highly-specific marker for exosomes and is not enriched in the larger microvesicle class of EVs.

## Introduction

Under normal conditions, mammalian cells secrete two size classes of extracellular vesicles (EVs), the exosomes and other small EVs (sEVs) that are <200 nm in diameter, and the microvesicles and other large EVs (lEVs) that are >200 nm in diameter(1–7). Of these, the exosomes are the most intriguing(1,3,7), as these sEVs are highly enriched in specific subsets of exosome cargo proteins, especially the exosomal tetraspanins (CD81, CD9, and CD63(5,8)), and scaffold proteins (e.g. syntenin(9,10)). In contrast, microvesicles and other lEVs have a composition more similar to that of the cell and do not appear to be enriched in any particular set of proteins or RNAs(1,7). In addition to the EVs released by healthy cells, an even wider array of EVs are released by infected, dying, dead, and damaged cells. These include the exosome-based enveloped viruses(11–14) and virus-like particles(11,15- 17), non-exosomal enveloped viruses, apoptotic blebs(18), the lEVs produced by cells infected with ‘non-enveloped viruses’ that are loaded with their virions(19,20), and mechanically-generated microsomes of plasma and other membranes(21).

Within the array of EVs that are produced by mammalian cells, exosomes are the most interesting and most abundant, transmitting signals, molecules, and nutrients to other cells, both nearby and far away(22–35), modifying the extracellular environment (matrices, interstitial and other biofluids(36–39)), and facilitating protein quality control in exosome-producing cells(40–43). Not surprisingly, exosomes contribute to numerous physiological and disease processes, are abundant in all biofluids, and hold high potential as carriers of vaccines and therapeutics(1,3). However, most of what we know about exosome biogenesis, composition, and function has come from *in vitro* studies. This has left unanswered many questions about how exosomes are made by different cell types *in vivo*; the extent to which different cell types contribute to the exosomes found in blood, lymph, and other biofluids; the amount and specificity of exosome traffic between different cell types, tissues, organs and organ systems; and how these processes change in response to different states of health and disease.

Exosomes can be identified by their combination of size (∼30-150 nm), topology (outside out, cytoplasm in), and the presence of highly-enriched exosome marker proteins. The most important of these are the tetraspanins CD81(5,6,8,11,16), CD9(9), and CD63(8,11,16), as they appear to be the most highly enriched of all exosome cargo proteins(5,44). Of these, CD81 is the most highly enriched exosome cargo protein known, and is loaded into exosomes ∼3-fold more efficiently than CD9 and ∼15-fold more efficiently than CD63(5). Furthermore, CD81 shows no enrichment in the larger microvesicles/lEVs(8), and is therefore a relatively specific marker of exosomes. We describe here a transgenic mouse model of exosome biogenesis that takes advantage of CD81’s unique properties as an exosome tracer protein, and validate the utility of this *exomap1* mouse for *in vivo* studies of exosome biology, including the cell type-specific contributions to the blood exosome population.

## METHODS

### Nucleic acids

The plasmid pFF077 was assembled by ligating synthetic DNAs carrying the Amp gene, bacterial origin of replication, attB1 segment(45), the CAG enhancer/promoter(46), a codon-optimized ORF that encodes MTS-tdTomato(47) flanked by two loxP511 sites(48), the HsCD81mNG ORF from pJM1358(49), the WPRE element(50), the bGH polyadenylation signal(51) and the attB2 segment(45). The MTS-tdTomato ORF was codon-optimized to minimize recombination between the direct DNA repeats present in the original tdTomato ORF(47). The plasmid pJM776 expresses iCre, a codon optimized form of Cre recombinase(52). The mRNA encoding ΦC31 integrase was synthesized as previously described(45).

### Cell culture and transfection

HEK293 cells were obtained from ATCC (CRL-1573) and grown in complete media (CM; DMEM, 10% fetal bovine serum, 1% penicillin/streptomycin). HEK293 cells were transfected with lipofectamine 3000 according to the manufacturer’s instructions (Thermo).

### Fluorescence microscopy of HEK293 cells

HEK293 cells transfected with either pFF077 or co-transfected with pFF077 and pJM776 were seeded onto sterile, poly-D-lysine-coated cover glasses, grown overnight, then fixed (3.7% formaldehyde in PBS for 30 minutes), permeabilized (1% Triton X-100 in PBS for 5 minutes), stained with DAPI, washed, and then mounted on glass slides. The cells were examined using BH2-RFCA microscope (Olympus) equipped with an Olympus S-plan Apo 63× 0.40 oil objective and a Sensicam QE (Cooke) digital camera using IPLab 3.6.3 software (Scanalytics, Inc.). Images were processed in Adobe Photoshop and figures were assembled in Adobe Illustrator.

### Preparation of sEV/exosomes from tissue culture supernatants

The analysis of exosomes released by HEK293 cells was carried out by culturing transfected HEK293 cells for three days, after which the conditioned media was collected and the cells were lysed in SDS-PAGE sample buffer. Exosomes were purified from conditioned media as previously described, with contaminating cells and cell debris removed by centrifugation at 5000 x g for 5 minutes, followed by filtration through a 200 nm pore diameter size sterile filtration unit. The resulting clarified tissue culture supernatant was then spun at 10,000 x *g* for 30 minutes, twice, to ensure removal of any residual large entities, and then spun at 100,000 x *g* for 2 hours to pellet sEVs. Exosome pellets were then resuspended in SDS-PAGE sample buffer, and cell and sEV lysates were stored at -80°C for later analysis.

### Immunoblots

Protein samples were generated by boiling cell and exosome samples in SDS-PAGE sample buffer. Mouse tissue proteins were generated by harvesting tissues of interest (liver, kidney, brain), mincing with a razor blade in PBS in a sterile tissue culture plate. Cells were then pelleted in a 2 mL microfuge tube and resuspended in 200 μL RIPA buffer (ThermoFisher Scientific, Cat. No. 89900, Waltham, MA, USA) and 2μL of Phosphatase/Protease inhibitor cocktail (ThermoFisher Scientific 78440). One stainless steel bead (5mm) was added to each tube, the tubes were inserted into a TissueLyser (Qiagen), and agitated at 50 vibrations/min until the tissue was homogenized properly. Insoluble material and the bead were removed by low-speed centrifugation for 1 min., and the resulting supernatant was clarified by centrifugation at 12,000 x *g* for 10 min at 4°C. The resulting cell lysate supernatants were transferred to new tubes and stored -80°C.

Protein samples in 1x SDS-PAGE sample buffer were separated by SDS-PAGE and electrophoretically transferred to PVDF membranes. Membranes were blocked, incubated in blocking solution containing primary antibody overnight at 4°C, washed, and incubated with an HRP conjugate of secondary antibody for 2 hours, then washed again. Membranes were incubated with ECL detection reagent and imaged using a digital light capture device. Gel images processed in Adobe Photoshop and figures were assembled in Adobe Illustrator. Unlabeled and labeled antibodies specific for human CD81, mouse CD9, mouse CD81, as well as control rat, hamster, and human IgG were acquired from commercial sources (Biolegend).

### Mice, gDNA extraction, and PCR

All procedures were conducted in accordance with the NIH Guide for the Care and Use of Laboratory Animals and were approved by the Institutional Animal Care and Use Committees of each participating laboratory. Mice were maintained in environmentally controlled vivarium, with food and water provided *ad libitum*. Adult mice were generated from breeding colonies at the Johns Hopkins University (*exomap1* founder line), Charles River laboratories (*exomap1* stock line), Massachusetts General Hospital, University of Kansas Medical Center, University of California Irvine, and University of California San Francisco/Veterans Administration San Francisco.

Pronuclear injection of H11P3 zygotes with linearized pFF077 DNA and ΦC31 integrase mRNA and injected zygotes were implanted into pseudo-pregnant females as previously described (the H11P3 mouse has three copies of attP in the mouse Hipp11 (H11) locus(45)). Pups were assessed for transgene integration by PCR analysis of tail snip genomic DNA using primers specific for the transgene and for sequences specific for the engineered H11P3 locus(45). Primers flanking the three attP sites in the H11P3 locus and internal to the *exomap1* transgene revealed that the MT124 founder mouse carried a single insertion of the *exomap1* transgene between attP sites 2 and 3 of the H11P3 locus. The MT124 mouse line was established by backcrossing the original *exomap1*^+/^ founder mouse to C57Bl/6 mice six times, with transgene carrier status monitored by fluorescence imaging of mice or mouse tail snips, and by PCR analysis of mouse tail snip gDNA using primers flanking the 5’ transgene insertion junction (5’- GGTGATAGGTGGCAAGTGGTATTCCG TAAG-3’ and 5’- CATATATGGGCTATGAACTAATGACC CCGT-3’, which yields a 447 bp-long DNA fragment) and/or the 3’ transgene insertion junction (5’-GCATCGCATTGTCTGAGTAGGTGTCA-3’) and 5’- CCGCGAAGTTCCTATACCTTTTG-3’, which yields a 301 bp-long DNA fragment). The extraction of gDNA from mouse tail snips and their analysis by PCR was performed using standard procedures.

Cre driver mice used in this study included the *Zp3*-Cre line C57BL/6-Tg(Zp3- cre)93Knw/J (Jackson Laboratories stock #003651(53)), the *Aromatase*-Cre line *cyp19a1*-Cre (generous gift from Dr. Jan A. Gossen(54), the *LysM*-Cre line B6.129P2- Lyz2tm1(cre)Ifo/J (Jackson Laboratories stock #004781(55)), the *HSA*-MCM line Tg(ACTA1-cre/Esr1*)2Kesr/J (Jackson Laboratories stock #025750(56)), the *Dat*- Cre line Slc6a3tm1(cre)Xz/J (Jackson Laboratories stock #020080(57)), and the *Camk2a*-Cre line B6.Cg-Tg(Camk2a- cre)T29-1Stl/J (Jackson Laboratories stock #005359(58)). Double transgenic F1 mice were obtained by crossing hemizygous Cre driver lines with hemizygous *exomap1^+/^* or homozygous *exomap1^+/+^* mice, with the presence of the *exomap1* transgene determined by fluorescence illumination of mice or of mouse tail snips and/or by gDNA PCR analysis, while the presence of Cre driver transgenes were assessed by gDNA PCR analysis.

### Virus infections

The AAV5/*Rpe65*-Cre virus was obtained from Vigene Biosciences (pAAV-Rpe65- Cre, AAV5 serotype, titer >1x10^13-10^14 gene copies/mL). Subjects were anesthetized with a 1-3% isoflurane/oxygen mixture and positioned in a Kopf stereotaxic frame with the incisor bar set to the flat-skull position. Brain microinjections were administered into the dorsal third ventricle at a volume of 2 μL at a rate of 0.1μl per minute across 20 minutes, and the injector remained in place for an additional 5 minutes to allow for diffusion. The coordinates were as follows: AP -1.00 mm, ML: midline at bregma, DV - 2.07 mm. The mice were sacrificed at least 2.5 after injection. Mice were then perfused with 4% paraformaldehyde, and organs were removed and placed in 30% sucrose. At least 72 hrs thereafter, tissue was sectioned on a cryostat, mounted onto microscope slides, coverslipped with Vectashield mounting media containing DAPI, and then examined by fluorescence microscopy. n=3 subjects were examined to verify expression patterns.

The AAV8/*TBG*-Cre viruses were obtained from commercial suppliers (1x10^13 genome copies (GC)/mL, CV17191-AV8, Charles River; or 2x10^13 GC/mL, 107787-AAV8, Addgene), diluted 1:20 into a final volume of 100 uL PBS, and delivered by intravenous injection into either the tail vein or retro-orbital sinus. Animals were maintained for 2- 4 weeks post-injection, after which tissue and blood samples were collected.

### Fluorescence microscopy of mouse cells and tissue sections

For the analysis of oocytes and ovarian tissues, mature female *exomap1*::*Zp3*-Cre and *exomap1*::*Cyp19a1*-Cre mice were sacrificed and ovaries were removed. Follicles were mechanically ruptured to disperse oocytes and somatic follicle (granulosa/cumulus) cells, and the dispersed cells were examined by fluorescence microscopy.

For fluorescence microscopy of tissue sections, mice were euthanized and then perfused with 4% paraformaldehyde. Organs were removed and placed in 30% sucrose. 3 days later, tissues were sectioned on a cryostat at 35 mm thickness, directly mounted onto microscope slides, then coverslipped with Vectashield with DAPI (Vector Labs, Cat# H-1200) and examined by fluorescence microscopy.

### Immunohistochemistry

Mice were euthanized, perfused with 4% paraformaldehyde, and organs were removed and placed in 30% sucrose. Three days later the tissues were frozen, sectioned (4 um) using a cryostat CM1850 (Leica, CA), mounted onto microscope slides, incubated with anti-HsCD81 monoclonal antibody linked to horseradish peroxidase, washed, and then processed for HRP immunohistochemistry.

### Blood, plasma, serum, and CSF collection

Blood samples were isolated by either retro- orbitual bleeding using Micro-hematocrit capillary tubes (Fisher Scientific), submandibular bleeding, or cardiac puncture into EDTA. Plasma samples were generated from bloods by immediately spinning EDTA bloods (within 60 minutes of collection) at 2000 x *g* for 10 minutes, followed by a second spin at 3000 x *g* for 10 minutes. Plasma samples were stored at -80°C. To isolate leukocytes, blood cells were collected by low-speed centrifugation, resuspended and incubated in ACK RBC lysis buffer according to the manufacturer’s instructions (Thermo), followed by pelleting of leukocytes by low-speed centrifugation and resuspension once again in ACK RBC lysis buffer. Leukocytes were then once again pelleted by low-speed centrifugation, after which they were interrogated by flow cytometry. Serum samples collected by cardiac puncture were allow to clot for 20-30 minutes and then centrifuged at 3000 x g for 10 minutes.

CSF samples were isolated from anesthetized mice as described(59). Briefly, the mouse’s head was fixed to a stereotaxic frame in the position that allowed the maximum opening of the cisterna magna. A Hamilton syringe (The Hamilton Company, Boston MA, USA) equipped with a 26-gauge needle or pulled glass pipette tip was inserted stereotactically in parallel to the brain axis to reach the cistern. The clear CSF (approximately 10 µL) was then drawn slowly to minimize possible contamination by blood or tissue.

### Flow cytometry

Flow cytometry of circulating leukocytes was performed by first blocking Fc receptors by incubating cells with TruStain FcX PLUS (anti-mouse CD16/32, Biolegend) for 10 minutes. Cell were then incubated with anti- CD45 & anti-CD11b (Biolegend), followed by purification of CD45+, CD11b+ cells on affinity capture beads. Fluorescence of MTS- tdTomato and HsCD81mNG in the CD45+, CD11b+ cell population was determined by flow cytometry using a Beckman Coulter CytoFLEX S cytometer. Histograms of fluorescence intensity were processed in Adobe Photoshop and figures were assembled in Adobe Illustrator.

### Isolation of exosomes/sEVs

Plasma and serum exosomes/sEVs were isolated by size exclusion column chromatography using pre-packed qEV columns (Izon), followed by concentration of peak exosome fractions by centrifugal flow filtration across a 10 kDa pore diameter filter, as recommended by the manufacturer (Millipore).

### Nanoparticle tracking analysis (NTA)

*Exomap1*::*Dat*-Cre male and female mice were used to obtain samples of CSF (n=15). CSF was first filtered with a 200 nm pore diameter cellulose acetate filter (Costar, RNase/DNase free, Cat#8161) and spun for 30 sec. Samples were then diluted 1:20 with nanopure H2O. The size distribution pattern of CSF sEVs was quantified with a NanoSight NS300 and nanoparticle tracking analysis (NTA) software version 3.3 with sCMOS camera and either (***i***) no filter (brightfield) to examine all EVs in the sample, or (***ii***) a green light filter to detect EVs with HsCD81mNG fluorescence. The camera settings were as follows: level 6, slider shutter 86, slider gain 15, 25.0 frames per second, number of frames 1498, temperature 21.7°C, syringe volume 0.6 - 1 mL, and syringe pump speed 100.

### Quantitative single molecule localization and total internal reflection fluorescence microscopy

25 mm diameter coverslips #1.5H (Thermo Fischer Scientific, Cat# NC9560650; Waltham, MA, USA) were functionalized with N-hydroxysuccinamide (NHS) groups, followed by covalent attachment of monoclonal antibodies that bind to epitopes in the ectodomain of either the mouse CD81 or human CD81 proteins. Raw biofluid samples from one *exomap1*::*Camk2a*-Cre mouse, one *exomap1*^+/^ mouse infected with AAV8/*TBG*-Cre virus, and one control *exomap1*^+/^ mouse were diluted in PBS to a final volume of 50 μL and placed on the surface of these antibody-coated coverslips at room temperature overnight in a humidified chamber. For the CSF samples, 1 μL of CSF was used for incubation with the anti-mouse CD81-functionalized coverslip and 6 μL of CSF were used for incubation with the anti-human CD81-functionalized coverslip. For the plasma samples, 1 μL of plasma was used for incubation with the anti-mouse CD81- functionalized coverslip and 5 μL of plasma were used for incubation with the anti-human CD81-functionalized coverslip. These coverslips were then washed with PBS containing 0.025% Tween 20 and EVs were labeled with one of the following affinity reagents: 1) a mixture of AF647-labeled antibodies specific for mouse CD9 (Biolegend, Cat. No. 124802, San Diego, CA, USA), mouse CD63 (Biolegend, Cat. No. 143902, San Diego, CA, USA), mouse CD81 (Biolegend, Cat. No. 104902, San Diego, CA, USA), and human CD81(Biolegend, Cat. No. 349502, San Diego, CA, USA) (*exomap1*::*Camk2a*-Cre mouse and control *exomap1*^+/^ mouse); 2) a mixture of AF647-labeled antibodies specific for mouse CD9, mouse CD63, and mouse CD81 (*exomap1*^+/^ mouse infected with AAV8/*TBG*-Cre virus and control *exomap1*^+/^ mouse); 3) AF647-labeled antibodies specific for human CD81 (*exomap1*^+/^ mouse infected with AAV8/*TBG*-Cre virus and control *exomap1*^+/^ mouse). All antibodies were fluorescently labeled as described previously at a molar ratio of ∼1(60)). Samples were fixed and stored as described previously(61,62).

For imaging, coverslips were placed in Attofluor cell chambers (Thermo Fisher Scientific) loaded with direct stochastic optical reconstruction microscopy (dSTORM) imaging buffer(63). N-STORM super-resolution microscope (Nikon Instruments; Melville, NY, USA) was used for TIRF and SMLM imaging using 488 nm and 640 nm lasers, respectively using microscope components described previously(64). Images (10 regions of interest (ROI) for CSF samples and 20 ROI for plasma samples) were acquired using NIS-Elements software (Nikon Instruments). SMLM images were processed using N-STORM Offline Analysis Module of the NIS-Elements software to localize peaks. The data were analyzed with Matlab R2022a (MathWorks; Natick, MA, USA) using the Voronoi tessellation algorithm(61,62) with a minimum of 40 localization points per cluster and an average of 14 localizations per single fluorescent tetraspanin antibody.

### SPIR-IFM imaging

Custom SPIR imaging chips derivatized with antibodies specific for mouse CD81, mouse CD9, and human CD81, and with control IgG from rat, hamster, and human, were generated by Nanoview (Waltham, MA). Serum samples were collected from control mice, *exomap1*+/ mice, and *exomap1*::AAV8/*TBG*-Cre mice (4 days post-infection) and sEVs were collected by filtration and SEC. sEVs/exosomes were then incubated on the SPIR imaging chips overnight at RT, washed, incubated with Alexa Fluor 555-labeled anti-human CD81, washed again, and interrogated by interferometric reflectance imaging to detect exosomes/sEVs, and by conventional fluorescence microscopy to measure vesicle-associated (***i***) HsCD81mNG fluorescence and (***ii***) Alexa Fluor 555 fluorescence.

## Results

### Design and validation of the exomap1 transgene

To design a useful transgenic mouse model of exosome biology, we first identified four ideal properties of its exosome marker/tracer protein. *First*, this marker protein should be based on an intact exosome protein that is highly enriched in exosomes, for the simple reason that enrichment is the most rigorous sign of active biogenesis. *Second*, the exosome cargo protein should be an integral membrane protein to preclude its release into biofluids as free protein, a problem that plagues many non-membrane proteins that have been erroneously referred to as exosome markers. *Third*, the cargo protein should be free of adverse effects on cell physiology and animal health. *Fourth*, it should have an extracellular epitope to which specific monoclonal antibodies are available, so that the marked exosomes can be selectively immunopurified and characterized. *Fifth*, the exosome marker protein should be expressed as a fusion protein to a bright fluorescent protein, so that its expression in target cells can be validated by fluorescence microscopy.

These considerations led us to select HsCD81mNG as the exosome marker/tracer protein in *exomap1* mice, as HsCD81 is: (***i***) the most highly-enriched exosomal protein yet described(5,8,65); (***ii***) not enriched in the larger microvesicle class of EVs (8); (***iii***) an integral membrane protein tetraspanin with four transmembrane domains, and is therefore incapable of being secreted as a soluble, non-vesicular protein; (***iv***) not associated with adverse effects on animal health; (***v***) detectable by species-specific monoclonal antibodies that bind extracellular epitopes in the second extracellular loop of HsCD81 but not mouse CD81; and (***vi***) efficiently loaded into exosomes when fused at its C-terminus to the N-terminus of the bright green fluorescent protein mNeonGreen (HsCD81mNG)(66).

These cargo-specific considerations were combined with general standards of mouse transgene design, which have validated the utility of (***vii***) inserting transgenes in safe harbor loci of the mouse genome such as the H11 locus (45), (***viii***) the CAG promoter/enhancer element to drive strong transgene expression in most mouse cells(67), (***ix***) the use of Cre recombinase and loxP site-based recombination to trigger transgene expression in specific cell types and/or at specific times(68), and (***x***) engineering the transgene to express a second fluorescent reporter prior to Cre-mediated transgene recombination(69).

We therefore assembled the *exomap1* transgene transfer vector pFF077 as a ΦC31 integrase donor vector(70) that carries a Cre-regulated transgene flanked by attB sites (***Fig. 1A***), so that the transgene could in the future be inserted into attP sites of the H11P3 mouse line (the H11P3 mouse line carries three attP sites integrated at the Hip11 locus(45)). Between these attB sites, the *exomap1* transgene consists of the CAG promoter, a loxP site, an open reading frame (ORF) encoding the mitochondrial red fluorescent protein MTS-tdTomato, multiple stop codons in each reading frame, a second loxP site, the HsCD81mNG ORF(49), and a 3’ untranslated region containing both the woodchuck hepatitis virus post-transcriptional regulatory element (WPRE)(50) and the polyadenylation signal from the bovine growth hormone gene (bGH).

**Figure 1.**
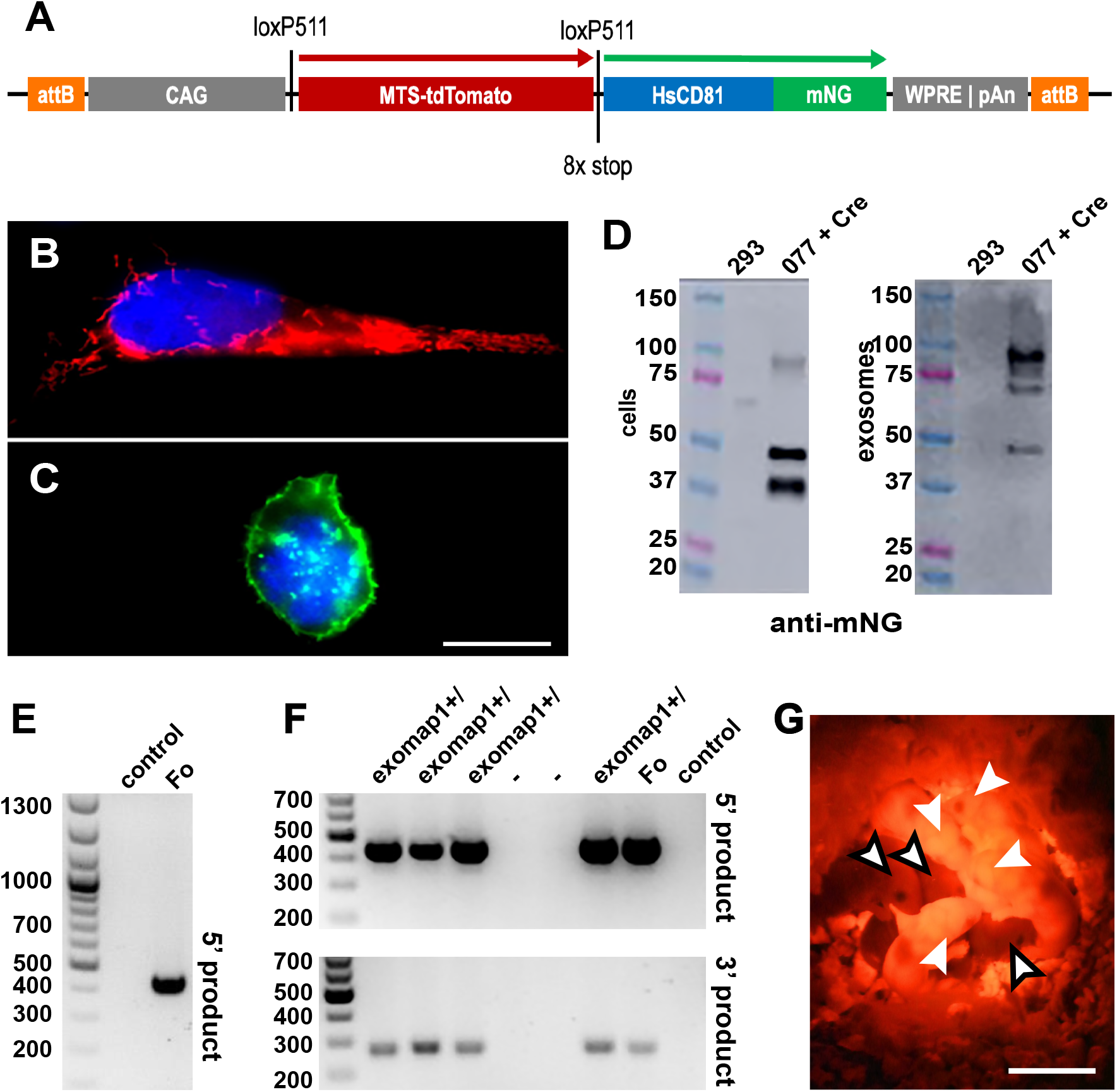
Design, genetics, and red fluorescence of exomap1^+/-^ mice. (**A**) Line diagram of the *exomap1* transfer vector. (**B**, **C**) Fluorescence micrographs of DAPI-stained HEK293 cells transfected with (B) pFF077 or (C) pFF077 + pJM776 that had been fixed and stained with DAPI. Bar, 10 um. (**D**) Anti-CD81 immunoblot of cell and exosome fractions of (left lanes) HEK293 cells and (right lanes) HEK293 cells expressing HsCD81mNG. MW size standards are in kDa. (**E**) Ethidium bromide-stained agarose gel electropherogram of genomic DNA (gDNA) PCR products generated using gDNAs extracted from (control) non-transgenic mouse and the (Fo) *exomap1* founder mouse using the H11 locus 5’ primer and the CAG promoter 3’ primer, showing the 447 bp product that is diagnostic for the *exomap1* transgene. MW size markers are in bp. (**F**) Ethidium bromide-stained agarose gel electropherograms of PCR reaction carried out with gDNAs extracted from the tails of six F1 progeny from a cross between the (Fo) founder mouse and a non-transgenic control mouse, as well as gDNAs from the founder mouse and a non-transgenic control mouse. Upper panel shows products obtained using the H11 locus 5’ primer and the CAG promoter 3’ primer. Lower panel shows products obtained using the bGH-pA 5’ primer and the FRTR2 3’ primer, with an expected *exomap1* transgene-specific product of 311 bp. MW size markers are in bp. (**G** Fluorescence micrograph of three F1 mice, illuminated with green light and imaged with a red filtered camera. White arrows point to heads to four *exomap1*^+/^ carrier mice, readily identifiable by their red fluorescent ears, feet, and tails. Black bordered arrows point to the heads of three non-transgenic littermates. Bar, 1 cm.

To ensure that the *exomap1* transgene in pFF077 was Cre responsive, we transfected it into HEK293 cells on its own or together with a Cre recombinase-expressing plasmid (pJM775). The next day, the transfected cells were seeded onto sterile cover glasses, and on day two the adhered cells were fixed, stained with DAPI, and examined by fluorescence microscopy. Cells transfected with pFF077 alone expressed bright red fluorescence in the cells’ mitochondria (***Fig. 1B***). In contrast, cells that had been co-transfected with pFF077 and pJM775 displayed bright green HsCD81mNG fluorescence, most of which was localized to the plasma membrane, though some was also detected at intracellular compartments (***Fig. 1C***).

Immunoblot analysis revealed that cells and exosomes contained multiple and distinct forms of HsCD81mNG (***Fig. 1D***). Specifically, we found most of the cell-associated HsCD81mNG migrated at its predicted molecular mass of ∼50 kDa but also at a breakdown size of ∼40 kDa. Cells also contained a small amount of HsCD81mNG that migrated at the expected size of an HsCD81mNG dimer, ∼100 kDa. HsCD81mNG proteins found in exosomes were a bit different, as most of the exosomal HsCD81mNG migrated at or slightly below the expected size of the HsCD81mNG homodimer, with the remaining immunoreactive protein migrating at the expected ∼50 kDa size of the HsCD81mNG monomer, and little if any of ∼40 kDa breakdown product that had been detected in cell lysates. The finding that exosomes were enriched for an SDS-resistant, higher molecular mass form of HsCD81mNG has some precedent, as we previously demonstrated that high order oligomerization facilitates the loading of plasma membrane proteins into exosomes(16).

### Genetics of the exomap1 mouse

To make the *exomap1* mouse, we linearized pFF077 by restriction enzyme cleavage at sites flanking the two attB sites, followed by purification of the resulting linear DNA donor fragment. This DNA was mixed with mRNA encoding the Φ131 integrase protein and injected into the pronucleus of zygotes of the H11P3 mouse. This transgenic mouse line carries three attP sites at the H11 safe harbor locus, and is designed to receive attB-flanked DNAs between the H11 locus attP1, attP2, and/or attP3 sites(45). Injected zygotes were then implanted into pseudo-pregnant females and the resulting pups were screened by PCR of genomic DNA (gDNA). The MT124 founder mouse resulted from insertion of the *exomap1* transgene between attP sites 2 and 3 of the H11P3 locus, and can be tracked through horizontal transmission by PCR reactions that flank the 5’ attP/B insertion site and the 3’ attB/P insertion site (***Fig. 1E, F***).

Hemizygous *exomap1*^+/^ mice express MTS- tdTomato throughout the body, allowing the optical identification of mice that carry the *exomap1* transgene upon proper illumination (***Fig. 1G***). Husbandry of *exomap1*^+/^ mice revealed that the *exomap1* transgene was inherited in a Mendelian pattern, with no evidence for adverse effects. Specifically, within a set of 21 crosses of homozygous *exomap1*^+/+^ mice (for the 29 months of 08/2020 to 01/2023), the average litter size was 6, the sex ratio was 1, and of the 10 animals that were kept for >270 days as long-term sources of the *exomap1* line, 5 were male and 5 were female.

### Oocyte-specific HsCD81mNG expression

To test whether Cre activates the mouse *exomap1*transgene, we first crossed *exomap1*^+/^ carriers to a *Zp3*-Cre mouse line(53). Zona pellucida 3 (*Zp3*) is expressed solely in the oocyte, with female F1 mice from the *exomap1*^+/^ x *Zp3*-Cre cross expected to express HsCD81mNG in the oocyte but not their surrounding cumulus cells. This prediction was borne out by fluorescence microscopy of oocytes and their surrounding cumulus cells, as oocytes displayed strong expression of HsCD81mNG and no substantial MTS-tdTomato fluorescence, while the somatic cumulus cells displayed strong MTS-tdTomato expression but no HsCD81mNG fluorescence (***Fig. 2A***). Close examination of these images confirmed that oocytes localized HsCD81mNG primarily to the plasma membrane, including the microvilli that protrude from the oocyte surface(71), though some of the fluorescence was also detected in internal compartments, possibly endosomes, lysosomes, and/or deep invaginations of the plasma membrane(72–74).

**Figure 2.**
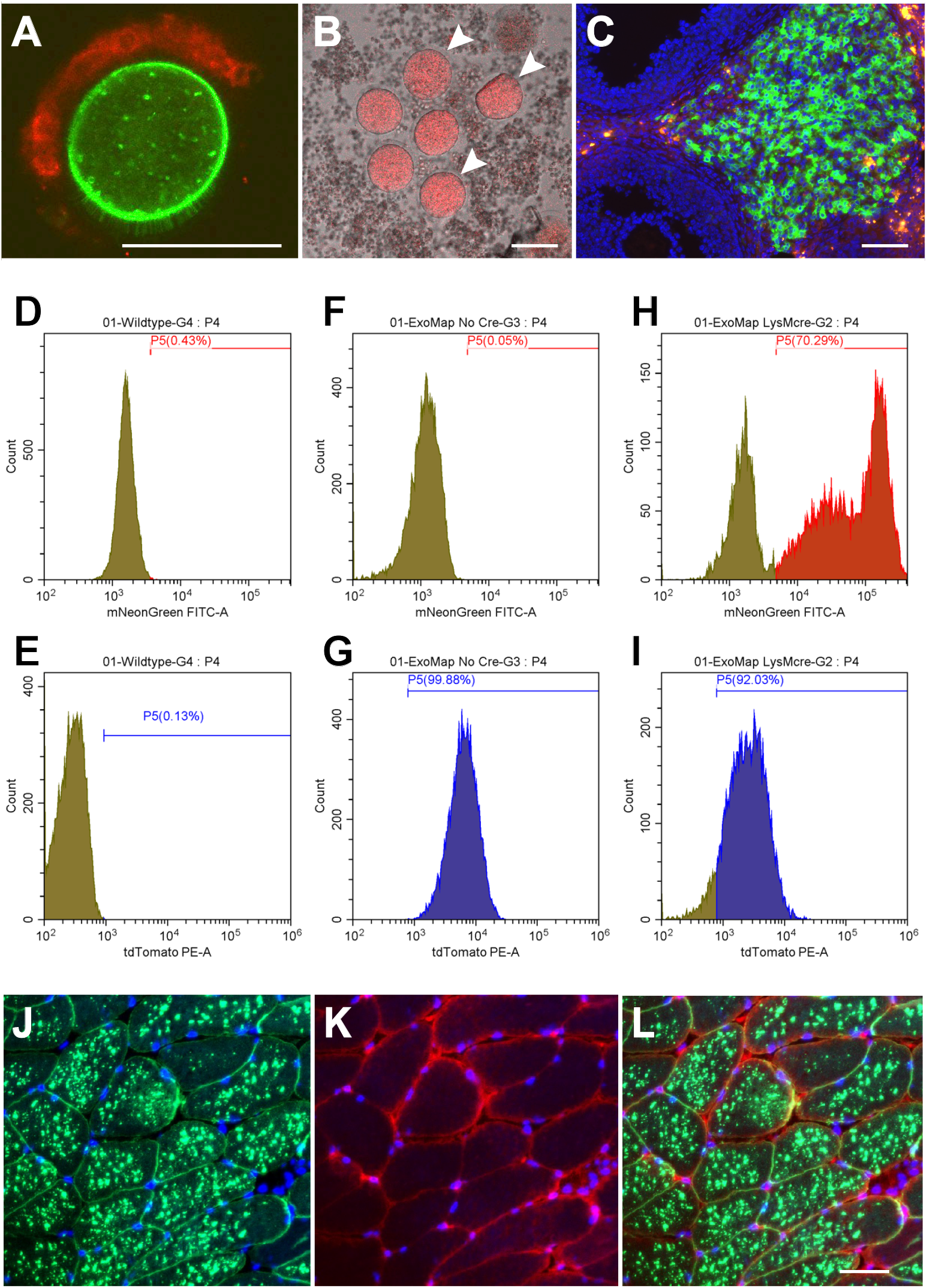
Cre-triggered expression of HsCD81mNG in exomap1 mice. (**A**) Fluorescence micrograph showing red and green fluorescence of an oocyte and a cluster of attached cumulus cells from an *exomap1*::*Zp3*-Cre mouse. Bar, 100 um (**B**) Fluorescence micrograph of oocytes and granulosa cells from *exomap1*::*Cyp19a1*-Cre mice showing lack of *exomap1* transgene expression in granulosa cells and strong exomap1 (MTS-tdTomato) expression in oocytes. White arrowheads point to oocytes. Bar, 100 um. (**C**) Fluorescence micrograph of DAPI-stained section through an *exomap1*::*Cyp19a1*-Cre mouse ovary, showing the absence of exomap1 transgene expression in granulosa cells of developing follicles (blue DAPI positive only) as well as strong *exomap1* transgene expression in corpus luteum cells, with most cells expressing HsCD81mNG while some expressed MTS-tdTomato and no HsCD81mNG. Bar, 100 um. (**D-I**) Flow cytometry histograms of CD45+, CD11b+ cells collected from the blood of (D, E) control mice, (F, G) *exomap1*^+/^ mice, and (H, I) *exomap1*::*LysM*-Cre animals showing plots of cell number vs brightness of (D, F, H) HsCD81mNG fluorescence and (E, G, I) MTS-tdTomato fluorescence. (**J-K**) Fluorescence micrographs of a cross section through the skeletal muscle of a tamoxifen-induced *exomap1*::*HSA*- MCM mouse stained with DAPI, sectioned, showing the images for (J) HsCD81mNG fluorescence and DAPI, (K) MTS-tdTomato fluorescence and DAPI. (L HsCD81mNG, MTS-tdTomato, and DAPI. Bar, 100 um.

### Complex expression of the exomap1 transgene in exomap1::Cyp19a1-Cre mice

Ovarian maturation is under strict hormonal control, with the expression of aromatase (encoded by the *Cyp19A1* gene) in somatic granulosa cells, allowing these cells to convert androgens into the intrafollicular estrogens necessary for oocyte maturation (75–77). Given the intriguing possibility that granulosa cell-derived exosomes might participate in follicular development, we attempted to activate HsCD81mNG in these cells by crossing *exomap1*^+/^ carriers with a *Cyp19a1*-Cre driver(54). However, when we collected follicles from *exomap1*::*Cyp19a1*- Cre female mice, released their oocytes and granulosa cells by mechanical disruption, and examined the dispersed cells by fluorescence microscopy, we observed that the granulosa cells lacked HsCD81mNG fluorescence, and also expressed little if any MTS-tdTomato fluorescence (***Fig. 2B***). This contrasted sharply with the bright red fluorescence of the oocytes, which are visible in this particular image (their cumulus cells are absent).

To explore this observation further, we examined ovary sections. As shown here, the somatic granulosa cells of immature follicles showed no expression of either MTS- tdTomato or HsCD81mNG, staining only for DAPI, while cells of a corpus luteum, which develops from the granulosa cells in the aftermath of oocyte maturation and release, displayed high expression from the *exomap1* transgene, with most cells expressing HsCD81mNG and some instead expressing MTS-tdTomato. Although we do not know why granulosa cells of immature follicles failed to express the *exomap1* transgene, the most likely explanation is that it reflects an unanticipated deficiency in granulosa cell expression from the H11 locus rand/or CAG promoter.

### Cre-induced expression of HsCD81mNG in myeloid cells

The observation that the *exomap1* transgene was not expressed in granulosa cells led us to next test its expression in several cell types of high interest to our consortium. These include cells of the myeloid lineage (monocytes, neutrophils, macrophages, etc.), which play key roles in cardiac health. We therefore crossed *exomap1*^+/^ carriers with *LysM*-Cre mice, which express Cre in CD45^+^, CD11b^+^ blood cells(55,78). Flow cytometry of CD45^+^, CD11b^+^ blood cells from control mice established the background levels of fluorescence in the green (HsCD81mNG) and red (MTS-tdTomato) channels (***Fig. 2D, E***), while flow cytometry of CD45^+^, CD11b^+^ blood cells from *exomap1*^+/^ animals showed that they express high levels of MTS- tdTomato fluorescence but no HsCD81mNG fluorescence (***Fig. 2F, G***). In contrast, flow cytometry of the same cell population isolated from *exomap1*::*LysM*-Cre showed strong HsCD81mNG fluorescence in ∼70% of cells (***Fig. 3H***), with the lack of expression in the other ∼30% likely due to incomplete penetrance of the *LysM*-Cre driver(78). Interesting, these flow cytometry experiments revealed the persistence of MTS-tdTomato fluorescence in all but ∼10% of cells (***Fig. 2I***), indicating that mitochondria-localized MTS-tdTomato fluorescence can stay fairly high even after its ORF has been deleted from the *exomap1* transgene.

**Figure 3.**
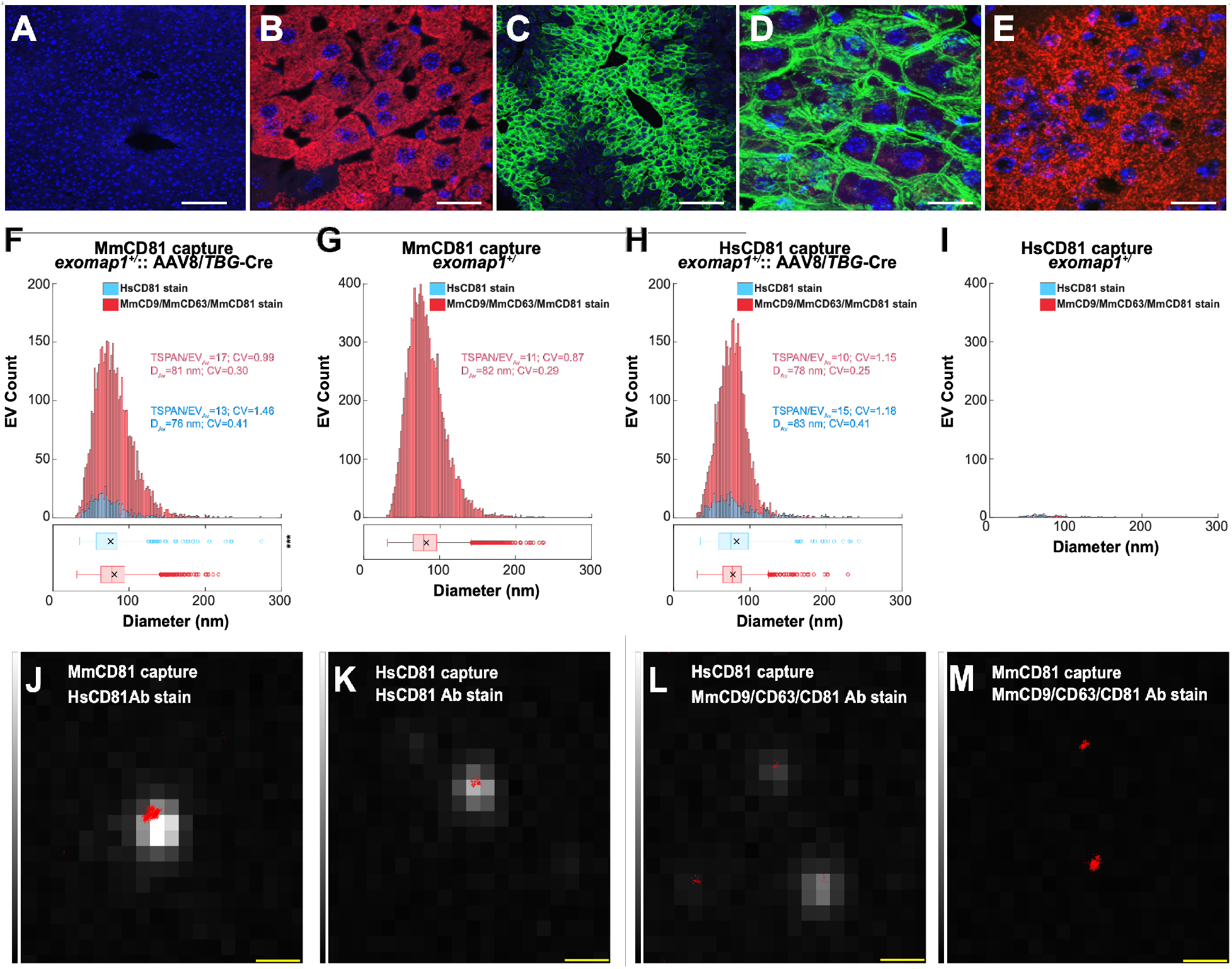
Hepatocytes load HsCD81mNG into mouse exosomes. (**A, B**) Fluorescence micrographs of liver sections from (A) a non-transgenic control mouse and (B) an *exomap1*^+/-^ mouse, stained with DAPI and imaged for (blue) DAPI and for (red) MTS-tdTomato fluorescence. Bar, 100 μm in A and 20 μm in B. (**C**) Fluorescence micrograph of a liver section obtained from an *exomap1*^+/^ mouse infected i.v. with AAV8/*TBG*-Cre virus, stained with DAPI, and imaged for (blue) DAPI and (green) HsCD81mNG fluorescence. Bar, 100 μm. (**D**) Fluorescence micrograph of a liver section obtained from an *exomap1*^+/^ mouse infected i.v. with AAV8/*TBG*-Cre virus, stained with DAPI, and imaged for (blue) DAPI, (green) HsCD81mNG fluorescence, and (red) MTS-tdTomato fluorescence. Bar, 20 μm. (**E**) Fluorescence micrograph of a brain section obtained from an *exomap1*^+/^ mouse infected i.v. with AAV8/*TBG*-Cre virus, stained with DAPI, and imaged for (blue) DAPI, (green) HsCD81mNG fluorescence. Bar, 20 nm. (**F-I**) Histograms of qSMLM data that we collected for raw plasma exosomes immunopurified on coverslips derivatized with antibodies specific for (F, G) MmCD81 or (H, I) HsCD81, from (F, H) an exomap1+/- mouse infected i.v. with AAV8/TBG-Cre virus and (G, I) an exomap1+/- mouse, that had been stained witheither (red) AF647-labeled anti-mouse tetraspanins (MmCD9, CD63, and CD81) or (blue) AF647-labeled anti-HsCD81. TSPAN/EV_Av_ denotes the average number of detected target markers per vesicle, CV denotes the coefficient of variation, and D_Av_ denotes the mean diameter. The box and whisker plot below histograms denote the mean (x), median (line), interquartile range (box), and point beyond 1.5-times the interquartile range (hollow dots). P values are included elsewhere (***Table S1***). (**J-M**) Fluorescence image overlays of (red dots) qSMLM localizations of bound surface tetraspanins and (greyscale) intrinsic HsCD81mNG fluorescence of raw plasma exosomes collected from an AAV8/*TBG*-Cre-infected *exomap1*^+/-^ mouse that had been affinity purified on (J, M) anti-MmCD81 or (K, L) anti-HsCD81 antibodies and stained with (J, K) AF647- labeled anti-HsCD81 antibody or (L, M) AF647-labeled antibodies to mouse exosomal tetraspanins. Bar, 500 nm.

### Skeletal muscle-specific HsCD81mNG expression

The biology of skeletal muscle exosomes are also of high interest, and we therefore crossed *exomap1*^+/^ mice with the tamoxifen-regulated human alpha-skeletal muscle actin (*HAS*)*-* Cre^ERT2^ mouse line(56). Adult *exomap1*::*HSA-*Cre^ERT2^ mice were injected with tamoxifen for 5 days, then ∼2-4 weeks later their muscle tissue was examined by fluorescence microscopy. These *exomap1*::*HSA-*Cre^ERT2^ mice displayed strong expression of HsCD81mNG in quadriceps muscle fibers (***Fig. 2J, L***), with MTS-tdTomato fluorescence emanating from the surrounding non-muscle cell types (e.g. pericytes, fibroblasts, adipocytes, endothelial cells, etc.) that border each muscle fiber (***Fig. 2K, L***). Lower resolution images confirmed the expression of MTS- tdTomato and absence of expression of HsCD81mNG in skeletal muscle of *exomap1*^+/^ mice, as well as the specific activation of HsCD81mNG expression in skeletal muscle of *exomap1*::*HSA-*Cre^ERT2^ mice (***fig. S1***).

### Hepatocyte expression of HsCD81mNG

We next induced the expression of HsCD81mNG in liver hepatocytes using an adeno-associated virus serotype 8 virus that carries a thyroxine binding globulin (TBG)- Cre transgene, AAV8/*TBG*-Cre, a virus that selectively activates Cre-dependent transgene expression in hepatocytes(79–83). The *exomap1* transgene is expressed well in hepatocytes, shown here by the MTS- tdTomato fluorescence in liver of *exomap1*^+/^ mice (***Fig. 3A, B***). When we infected *exomap1*^+/^ mice with AAV8/*TBG*-Cre, HsCD81mNG expression was induced in a large proportion of hepatocytes, especially hepatocytes that were proximal to liver sinusoids (***Fig. 3C***). Moreover, higher magnification images confirmed that hepatocytes localized HsCD81mNG primarily to the plasma membrane (***Fig. 3D***). This activation of HsCD81mNG expression was specific for liver, as brain tissue from AAV8/*TBG*-Cre-infected *exomap1*^+/^ mice expressed MTS-tdTomato, not HsCD81mNG (***Fig. 3E***).

### Mouse cells load HsCD81mNG into mouse exosomes

The hepatocyte expression of HsCD81mNG, together with the high likelihood that hepatocytes release large numbers of exosomes into the blood, allowed us to test whether mouse cells load HsCD81mNG protein into mouse exosomes *in vivo* by interrogating the immunophenotypes of blood exosomes collected from AAV8/*TBG*- Cre-infected *exomap1*^+/^ mice. Towards this end, we immunopurified exosomes from raw plasma samples using coverslips derivatized with monoclonal antibodies specific for either mouse CD81 (MmCD81), to bind all mouse exosomes, or HsCD81, to bind exosomes derived from HsCD81mNG- expressing cells. After overnight incubation, the coverslips were washed to remove unbound material and stained with either (***i***) a cocktail of Alexa Fluor 647 (AF647)-labeled monoclonal antibodies specific for the mouse exosome marker proteins MmCD9, MmCD63, and MmCD81, or (***ii***) an AF647- labeled monoclonal antibody specific for HsCD81. Each coverslip was then washed, fixed, and examined by quantitative single-molecule localization microscopy (qSMLM). These experiments revealed that a subset of mouse plasma exosomes from AAV8/*TBG*- Cre-infected *exomap1*^+/^ mice, immunopurified on a mouse-specific anti- MmCD81 antibody, stained positively with a human-specific anti-HsCD81 antibody (***Fig. 3F, G***). Furthermore, we found that HsCD81mNG-containing exosomes immunopurified on anti-HsCD81-specific antibodies also labeled positively with antibodies specific for the mouse forms of CD9, CD63, and/or CD81 (***Fig. 3H, I***). Together, these results provide compelling evidence that mouse cells load HsCD81mNG into vesicles that have a similar size (∼80 nm), topology (outside out), and composition (presence of mouse exosome markers) as mouse exosomes.

### Hepatocytes contribute ∼15% of blood exosomes

To estimate the hepatocyte contributions to the blood exosome populations, we quantified the percentage of HsCD81mNG positive exosomes present in the raw plasma of AAV8/*TBG*-Cre infected *exomap1*^+/^ mice. When we considered the number of exosomes captured on the anti-MmCD81 antibody that stained positively with the AF647-labeled anti-HsCD81 antibody (compared to staining with AF647-labeled anti-MmCD9/CD63/CD81 antibodies cocktail), the hepatocyte contribution to plasma exosomes appeared to be ∼11%. In particular, on average we detected HsCD8 on 23 exosomes per region of interest (ROI) and MmCD9/CD63/CD81 on 202 exosomes per ROI when results were normalized to 1uL of raw plasma (***Table 1***). As for the physical characteristics, those stained with anti- HsCD81 had an average diameter of 76 nm and an average of 13 detected HsCD81 molecules per immobilized exosome (***Fig. 3F, blue***). These values were highly similar to the exosomes stained with mouse exosome markers, which had an average diameter of 81 nm and an average of 17 detected mouse exosome markers (***Fig. 3F, red***). Importantly, the mouse CD81-positive exosomes collected from raw plasma of *exomap1*^+/^ mice stained robustly with the anti-mouse exosome marker antibodies. These exosomes had an average diameter 82 nm and an average of 11 detected mouse exosome marker proteins per vesicle (***Fig. 3G, red***). Only background staining (∼0.1%) was observed with the anti-HsCD81 antibody; this probe detected only 0.7 exosomes per ROI while the anti-mouse tetraspanin cocktail detected 517 exosomes per ROI when results were normalized to 1uL of raw plasma (***Table 1***). To ensure robust detection, in the preceding experiments, we required the presence of ∼ 3 detected marker proteins (equivalent to 40 localizations of a detection antibody across all acquired frames) before an exosome was counted as positive.

**Table 1.** Numbers of exosomes detected in plasma samples from *exomap1* and *exomap1*::AAV8/*TBG-Cre* mice. . Plasma was collected from *exomap1***^+/^**and *exomap1*::AAV8/*TBG*- Cre mice followed by immunopurification of sEVs on coverslips derivatized with anti-MmCD81 or anti-HsCD81 monoclonal antibodies. The immunopurified sEVs were then stained using a cocktail of fluorescently-tagged antibodies specific for mouse CD9, CD63, and CD81 or the anti- HsCD81 antibody only. Data are from 20 technical replicates of two independently-collected coverslips (values normalized to 1 uL of plasma).

|  |  | Mean | Median | SEM | CV |
| --- | --- | --- | --- | --- | --- |
| *exomap1::AAV8/TBG-Cre* mCD81 capture/mouse stain | EV/uL | 202 | 198 | 5.8 | 0.13 |
| *exomap1::AAV8/TBG-Cre* mCD81 capture/hCD81 stain | EV/uL | 23.0 | 22.0 | 1.4 | 0.28 |
| *exomap1::AAV8/TBG-Cre* hCD81 capture/mouse stain | EV/uL | 35.4 | 32.4 | 2.6 | 0.33 |
| *exomap1::AAV8/TBG-Cre* hCD81 capture/hCD81 stain | EV/uL | 4.9 | 5.1 | 0.4 | 0.33 |
| *exomap1*<sup>+/+</sup> mCD81 capture/mouse stain | EV/uL | 517 | 535 | 19.9 | 0.17 |
| *exomap1*<sup>+/+</sup> mCD81 capture/hCD81 stain | EV/uL | 0.7 | 0.0 | 0.2 | 1.16 |
| *exomap1*<sup>+/+</sup> hCD81 capture/mouse stain | EV/uL | 0.8 | 0.7 | 0.1 | 0.70 |
| *exomap1*<sup>+/+</sup> hCD81 capture/hCD81 stain | EV/uL | 0.5 | 0.3 | 0.1 | 1.13 |

The downside of this rigor is that it can underestimate the percentage of mouse exosomes that carry HsCD81. To avoid this underestimate, we compared the number of exosomes per ROI (results normalized to 1 uL of raw plasma), captured on either anti- HsCD81 antibodies or anti-MmCD81 antibodies; in both cases exosomes were stained with the cocktail of anti-mouse exosome markers. In this scenario, the capture of an HsCD81-positive exosome requires only a single HsCD81 molecule in the exosome membrane. From AAV8/*TBG*- Cre-infected *exomap1*^+/^ mice, we detected 35 exosomes per ROI on the anti-HsCD81 coverslips and 202 exosomes per ROI on the anti-MmCD81 coverslips (***Table 1***), yielding an estimate of the hepatocyte contribution to blood exosomes of ∼17%. As for the sizes of the mouse exosomes collected on anti- HsCD81 antibodies, they had a mean diameter of 78 nm and an average of 10 detected mouse exosome marker proteins (***Fig. 3H, red***), which was similar to those collected on anti-MmCD81 antibodies (***Fig. 3F, red***). As for the noise of our detection assay, our analysis of raw plasma from uninfected *exomap1*^+/^ mice indicates that it was only 0.2% (0.8/517 detected exosomes per ROI when results were normalized to 1 ul of raw plasma (***Fig. 3I, red***; ***Table 1***). Interestingly, only ∼5 exosomes per ROI collected on anti-HsCD81 antibodies stained positively with the anti-HsCD81 antibody (***Table 1***). These exosomes had a mean diameter of 83 nm and an average of 15 detected HsCD81 molecules, indicating this population was highly enriched in HsCD81 (***Fig. 3H, blue***). Our analysis of raw plasma from uninfected *exomap1*^+/^ mice indicates that 0.5 exosomes were detected per ROI when results were normalized to 1 ul of raw plasma (***Fig. 3I, blue***; ***Table 1***).

In addition to interrogating the exosomes from AAV8/TBG-Cre-infected exomap1+/ mice by qSMLM, we also assessed the intrinsic HsCD81mNG fluorescence by total internal reflection fluorescence (TIRF) microscopy. The results of these experiments added further evidence that HsCD81mNG is loaded into mouse exosomes. Specifically, we found that exosomes that stained positive with AF647-labeled anti-HsCD81 antibodies displayed a large number of EVs with high mNeonGreen (mNG) fluorescence (***Fig. 3J, K***), those captured on antiHsCD81 antibodies stained with antimouse exosomal markers were also likely to show at least some mNG fluorescence (***Fig. 3L***), while exosomes collected on anti-MmCD81 antibodies stained with anti-mouse exosomal markers were the least likely to display mNG fluorescence (***Fig. 3M***). Interrogation of plasma exosomes from AAV8/*TBG*-Cre-infected *exomap1*^+/^ mice by the coupled techniques of single-particle interferometric reflectance (SPIR) and immunofluorescence microscopy (IFM) also yielded supporting data that HsCD81mNG was loaded into exosomes (***Fig. S2***).

### HsCD81mNG activation in the brain

With these results in hand, we next tested whether exomap1 mouse could be used for studies of exosome biology in the central nervous system (CNS). A first test of this possibility was carried out by infecting *exomap1*^+/^ mice with AAV5/*Rpe65*-Cre, a virus designed to express Cre in a variety of neuronal support cells (astrocytes, microglia, etc.). This virus was delivered into *exomap1*^+/^ mice by intracerebroventricular (ICV) injection, followed three weeks later by fluorescence microscopy of brain tissue. This revealed the presence of HsCD81mNG fluorescence in individual cells in white matter tracks, including the corpus callosum (***Fig. 4A***), the posterior commissure (***Fig. 4B***), and in subventricular regions surrounding the lateral ventricle (***Fig. 4C***).

**Figure 4.**
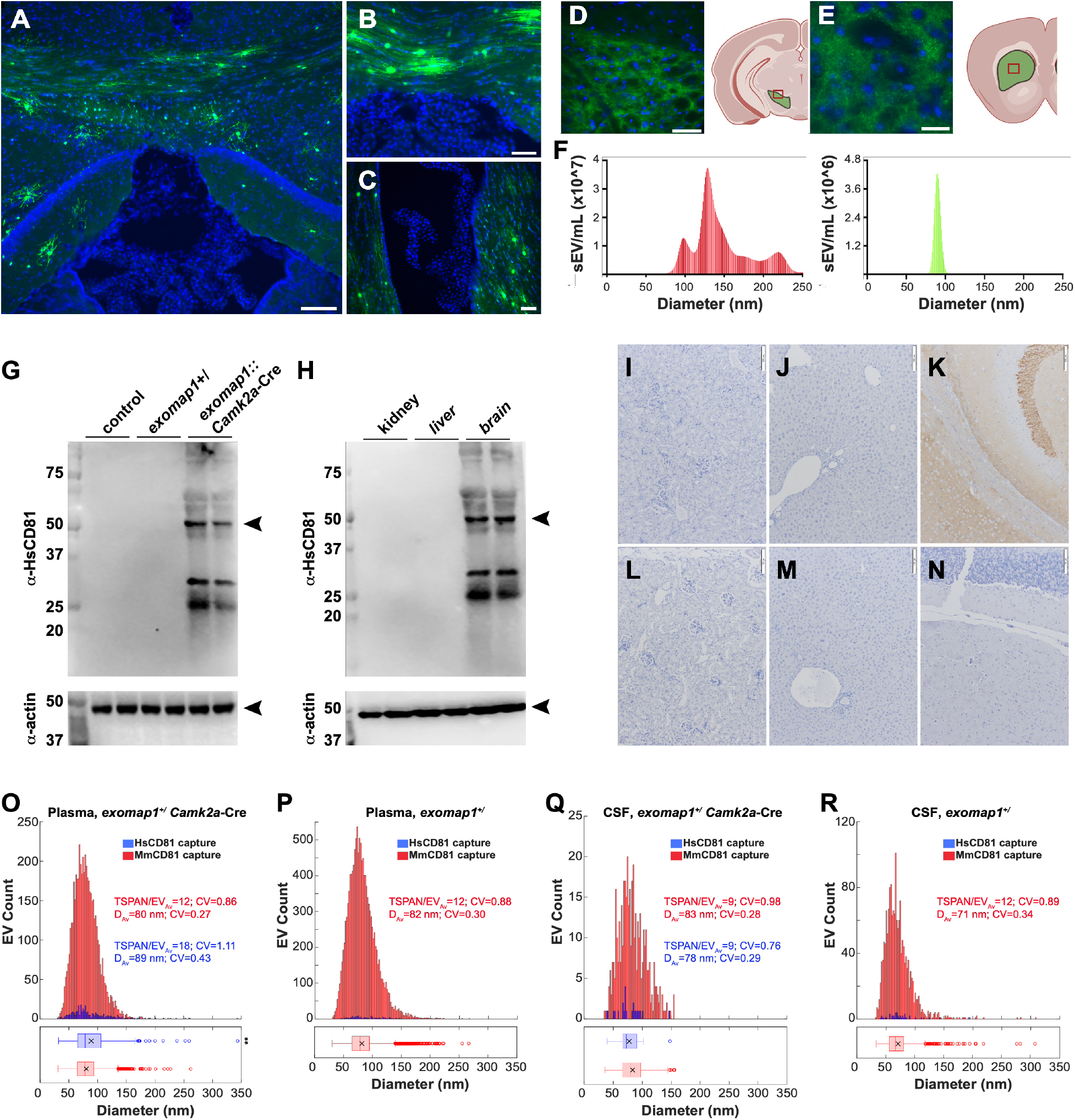
Activation of the exomap1 transgene in brain and neurons. (**A-C**) Blue/green fluorescence micrographs of DAPI-stained brain sections of *exomap1*^+/-^ mice injected with AAV5/*Rpe65*-Cre virus, showing HsCD81mNG fluorescence in cells of (**A**) the corpus callosum and hippocampus, (**B**) posterior commissure, and (**C**) lateral ventricle. Bar in A, 100 um; Bar in B and C, 50 um. (**D, E**) Fluorescence micrographs of DAPI-stained brain sections of *exomap1*::*Dat*- Cre mice showing HsCD81mNG fluorescence in (**D**) cells of the ventral tegmental area and (**E**) the axons in the striatum. Bar in A, 50 um; bar in B, 25 um. Cartoons depicting the brain location of these sections are presented to the right of each micrograph, and were generated in BioRender. (**F**) NTA histograms of raw CSF samples that had been passed through a 200 nm pore diameter size filter, showing (left histograpm) the size distribution profile of all CSF sEVs and (right histogram) the size distribution profile of mNeonGreen-positive sEVs. (**G**) Anti-HsCD81 immunoblot of brain protein extracts from control mice, *exomap1*^+/-^ mice, *exomap1*::*Camk2a*-Cre mice. MW size standards are in kDa. (**H**) Anti-HsCD81 immunoblot of kidney, liver, and brain protein extracts *exomap1*::*Camk2a*-Cre mice probed with antibodies specific for HsCD81. MW size standards are in kDa. (**I-N**) Micrographs showing anti-HsCD81immunohistochemical staining in tissue sections of (I, L) kidney, (J, M) liver, and (K, N) brain from (I-K) *exomap1*::*Camk2a*-Cre mice and (L-N) *exomap1*^+/^ mice. Bar, 100um. (**O, P**) Histograms of qSMLM data for raw plasma samples collected from (O) *exomap1*::*Camk2a*-Cre mouse or (P) *exomap1*^+/^ mouse, immunopurified on coverslips functionalized with (red) anti-MmCD81 antibody or (blue) anti- HsCD81 antibody, and stained with a cocktail of AF647-labeled antibodies to exosomal tetraspanins. TSPAN/EV_Av_ denotes the average number of detected target markers per vesicle, CV denotes the coefficient of variation, and D_Av_ denotes the mean diameter. The box and whisker plot below histograms denote the mean (x), median (line), interquartile range (box), and point beyond 1.5-times the interquartile range (hollowdots). (**Q, R**) Histograms of qSMLM data collected for raw CSF samples collected from (Q) *exomap1*::*Camk2a*-Cre mouse or (R) *exomap1*+/- mouse, immunopurified on coverslips functionalized with (red) anti-MmCD81 antibody or (blue) anti- HsCD81 antibody, and stained with a cocktail of AF647-labeled antibodies to exosomal tetraspanins. TSPAN/EV_Av_ denotes the average number of detected target markers per vesicle, CV denotes the coefficient of variation, and D_Av_ denotes the mean diameter. The box and whisker plot below histograms denote the mean (x), median (line), interquartile range (box), and point beyond 1.5-times the interquartile range (hollow dots). P values are included elsewhere (***Table S1***).

To test whether Cre expression can drive HsCD81mNG expression in CNS neurons, *exomap1*^+/^ mice were crossed with the dopamine transporter (*Dat*) Cre driver, *Dat*- Cre(57). *Dat* is expressed in dopamine neurons of the midbrain, and fluorescence microscopy of *exomap1*::*Dat*-Cre brain sections confirmed HsCD81mNG expression in cells of the ventral tegmental area (***Fig. 4D***) and axons in the striatum (***Fig. 4E***). Imaging for both HsCD81mNG and MTS- tdTomato fluorescence revealed the complexity of HsCD81mNG and MTS- tdTomato expression in the midbrain (***fig. S3***), including the presence of some cells that simply do not express high levels of either of these *exomap1* transgene products.

Given that the midbrain is proximal to the central aqueduct, we also tested whether HsCD81mNG fluorescence could be detected in the cerebrospinal fluid (CSF) of *exomap1*::*Dat*-Cre mice. For these experiments, CSF samples were collected from *exomap1*::*Dat*-Cre mice, passed through a 200 nm pore diameter filter, then examined by nanoparticle tracking analysis (NTA) coupled to fluorescent particle detection. This revealed the size distribution profile of sEVs in mouse CSF, and also identified those sEVs that displayed HsCD81mNG fluorescence above background. Importantly, HsCD81mNG fluorescence was detected in exosome-sized vesicles of ∼80-90 nm diameter (***Fig. 4F***). However, only ∼1% of the sEVs had detectable levels of HsCD81mNG fluorescence, indicating that Dat+ neurons contribute relatively few exosomes to the CSF.

### Camk2a neurons contribute <1% of blood exosomes

There is high interest in the detection of brain disease by liquid biopsy of blood EVs (84–88). To determine whether neuron-derived exosomes could be detected in the blood, we crossed *exomap1*^+/^ mice with a Cre driver line that expresses Cre recombinase under the control of the calcium/calmodulin-dependent protein kinase 2A (*Camk2a*) gene(58,89). Unlike *Dat*, which is expressed in a limited subset of neurons of the midbrain, *Camk2a*- Cre mice activate Cre-dependent transgenes in nearly all neurons, both central and peripheral. Immunoblot analysis confirmed the selective yet robust expression of HsCD81mNG in brain tissue from *exomap1*::*Camk2a*-Cre mice and its absence from control tissues (kidney and liver) (***Fig. 4G,H***), a result we also confirmed tissue immunohistochemistry (***Fig 4I-N***).

To measure the neuronal contributions to blood EV populations, we used qSMLM to interrogate raw plasma from *exomap1*^+/^ mice and *exomap1*::*Camk2a*-Cre mice. Specifically, raw plasma samples were incubated onto coverslips derivatized with antibodies specific for either MmCD81 or HsCD81, washed, stained with a cocktail of AF647-labeled antibodies specific for MmCD9, MmCD63, MmCD81, and HsCD81, then washed again, fixed, and examined by qSMLM. These experiments showed once again that exosomes captured with anti-MmCD81 antibodies were abundant in the plasma of both *exomap1*::*Camk2a*-Cre and *exomap1*^+/^ mice; they had average diameters of ∼80 nm (80 nm and 82 nm, respectively) and carried an average of 12 detected exosomal tetraspanins per sEV (***Fig. 4O, P, red***). Exosomes captured with anti-HsCD81 antibodies from plasma of *exomap1*::*Camk2a*-Cre mouse they had average diameters of ∼89 nm and carried an average of 18 detected exosomal tetraspanins per sEV (***Fig. 4O, blue***). The proportion of plasma exosomes that were derived from neurons of *exomap1*::*Camk2a*- Cre mice appeared to be low, only ∼1.3%, as we detected only 3.3 exosomes per ROI on our anti-HsCD81-capture coverslips compared to ∼259 exosomes per ROI on the anti-MmCD81 coverslips when results were normalized to 1uL of raw plasma (***Table 2***). Background in these experiments appeared to be ∼0.3%, as our analysis of raw plasma from *exomap1*^+/^ carriers detected 1.7 exosomes per ROI on our anti-HsCD81-capture coverslips compared to ∼648 exosomes per ROI on the antiMmCD81 coverslips when results were normalized to 1uL of raw plasma (***Table 2***).

**Table 2.** Numbers of exosomes detected in plasma samples from *exomap1* and *exomap1*::*camk2a*-Cre mice. . Plasma was collected from *exomap1***^+/^** and *exomap1*::*camk2a*-Cre mice, followed by immunopurification of sEVs on coverslips derivatized with anti-MmCD81 or anti-HsCD81 monoclonal antibodies. The immunopurified sEVs were then stained using a cocktail of fluorescently-tagged antibodies specific for mouse CD9, CD63, and CD81 and human CD81. Data are from 20 ROIs of two independently-imaged coverslips (values normalized to 1 uL of plasma).

|  |  | Mean | Median | SEM | CV |
| --- | --- | --- | --- | --- | --- |
| *exomap1::camk2a-Cre* mCD81 capture | EV/uL | 259 | 254 | 7.1 | 0.12 |
| *exomap1::camk2a-Cre* hCD81 capture | EV/uL | 3.3 | 3.2 | 0.2 | 0.28 |
| *exomap1*<sup>+/+</sup> mCD81 capture | EV/uL | 648 | 654 | 29 | 0.20 |
| *exomap1*<sup>+/+</sup> hCD81 capture | EV/uL | 1.7 | 1.3 | 0.2 | 0.60 |

To determine whether the pan-neuronal expression of HsCD81mNG led to a greater proportional contribution to CSF, we performed a parallel analysis of CSF collected from these mice. Exosomes were immunopurified from raw CSF by incubation on coverslips derivatized with anti-MmCD81 or anti-HsCD81 antibodies, stained with the cocktail of AF647-labeled antibodies specific for MmCD9, MmCD63, Mm CD81, and HsCD81, then washed, fixed, and examined by qSMLM. Exosomes captured from CSF of exomap1::Camk2a-Cre mouse had average diameters of ∼80 nm (83 nm and 78 nm, for capture with antibodies specific for either MmCD81 or HsCD81, respectively) and carried an average of 9 detected exosomal tetraspanins per sEV (***Fig. 4Q***). The proportion of CSF exosomes captured on anti-HsCD81-derivatized coverslips was ∼0.9% of those captured on antiMmCD81- derivatized coverslips (0.4/42 per ROI), compared to 0.4% for raw CSF from control *exomap1*^+/^ mice (0.7/161 per ROI); results were normalized to 1uL of CSF (***Table 3; Fig. 4Q, R***).

**Table 3.**
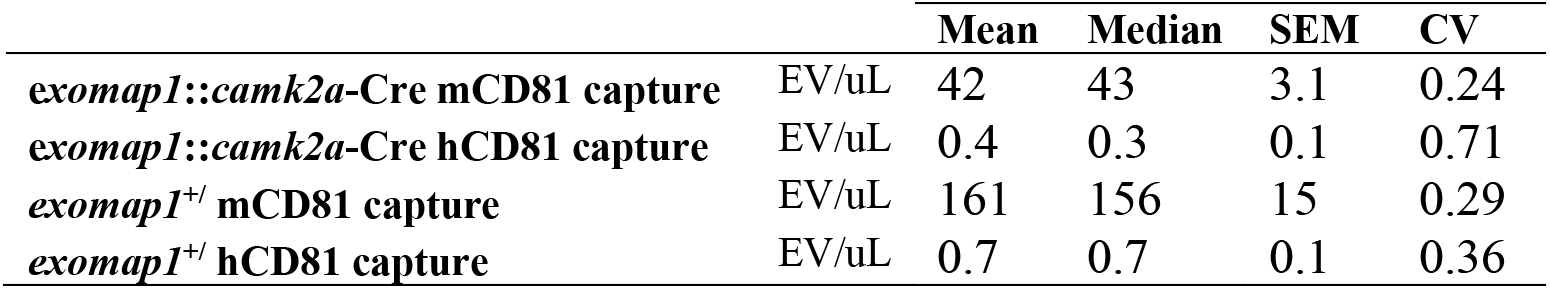
Numbers of exosomes detected in CSF samples from *exomap1* and *exomap1*::*camk2a*-Cre mice. CSF was collected from *exomap1***^+/^** and *exomap1*::*camk2a*-Cre mice followed by immunopurification of sEVs on coverslips derivatized with anti-MmCD81 or anti- HsCD81 monoclonal antibodies. The immunopurified sEVs were then stained using a cocktail of fluorescently-tagged antibodies specific for mouse CD9, CD63, and CD81 and human CD81. Data are from 10 ROIs (values normalized to 1 uL CSF).

## Discussion

The design of the *exomap1* mouse was based on an array of prior observations, including (***i***) the 25 years of data showing and affirming that CD81 is the most highly-enriched exosome cargo protein yet described(1,3,5,6,8,90), (***ii***) the lack of enrichment of CD81 in microvsicles/lEVs (***iii***) the fact that CD81 is an integral membrane tetraspanin and therefore never released from cells as free protein; (***iv***) the apparently benign effects of HsCD81mNG expression; (***v***) the commercial availability of monoclonal antibodies that are specific for the human and mouse forms of CD81; and (***vi***) the fact that mammalian cells correctly localize HsCD81mNG to the plasma membrane, secrete it from the cell in exosomes, and confer bright green fluorescence on expressing cells and exosomes(66). These were combined with generic advantages offered by (***vii***) transgene insertion at the H11 safe harbor locus(45), (***viii***) CAG-driven transgene expression(67), (***ix***) Cre-dependent expression of the exosome tracer protein(45), and (***x***) inclusion of a fluorescent reporter (MTS-tdTomato) to mark *exomap1*-expressing cells prior to Cre recombinase expression.

The data presented in this report indicate that each of these design elements worked largely as expected. Insertion of the CAG-driven *exomap1* transgene in the H11 locus created an *exomap1* mouse that expressed MTS- tdTomato in the vast majority of mouse cells and showed no expression of HsCD81mNG in the absence of Cre recombinase. Furthermore, the *exomap1* transgene was not associated with any deleterious traits that could be observed during the husbandry of this transgenic line as either hemizygous or homozygous carriers. The *exomap1* mouse also responded appropriately to Cre recombinase by activating the expression of HsCD81mNG in Cre-expressing cells, while also triggering a decline or loss of MTS- tdTomato expression in cell lines as diverse as oocytes, the corpus luteum, myeloid cells, skeletal muscle, hepatocytes, glia/astrocytes/microglia, and neurons.

### CD81 is a highly specific marker of exosomes

As noted above, the *exomap1* mouse was designed around the fact that HsCD81 is the most highly enriched exosome marker protein ever described. This observation was made first by Escola et al.(8) and confirmed again by us. In fact, CD81 is ∼15-fold more highly enriched in exosomes than the CD63(5,44,65). Escola et al. also reported that CD81 was largely absent from the larger microvesicle class of EV(8), and our unbiased analysis of CD81-containing EVs shows this once again. Specifically, when we immunopurified CD81-positive EVs directly from raw plasma and CSF samples, we found they had an average diameter of exosomes, ∼80 nm and that nearly all were under 200 nm in diameter. As for where these CD81- positive exosomes are made, the data presented here showed that diverse mouse cell types localized HsCD81 to the plasma membrane, consistent with our finding that the plasma membrane is the primary site of exosome cargo protein budding in various cell lines(5,16,44,65,91). Taken together, these and other results presented in this paper demonstrate that the *exomap1* mouse is a useful tool for *in vivo* studies of exosome biology.

### Hepatocytes and neurons contribute ∼15% and <1% of blood exosomes, respectively

In addition to validating the *exomap1* mouse as a useful model of exosome biogenesis, we demonstrated that the *exomap1* mouse can be used to estimate cell type-specific contributions to biofluid exosome populations. Specifically, we used qSMLM to immunophenotype individual exosomes that had been immunopurified directly from the raw plasma of mice that expressed HsCD81mNG in hepatocytes or neurons. These data indicate that hepatocytes contribute ∼15% of blood exosomes while neurons contribute <1% of blood exosomes. Although there may be mouse-to-mouse variations, these estimates correspond to the known vascular permeabilities of the endothelia that separate hepatocytes and neurons from the blood. Specifically, the liver sinusoidal endothelial cells possess numerous large transcellular pores (∼50-300 nm diameter)(92,93) that allow diffusion of large particles between hepatocytes and blood, whereas the blood-brain and blood- neuron barriers block the free flow of molecules larger than ∼1-5 nm(94). Furthermore, our estimate that hepatocytes contribute ∼15% of blood EVs is consistent with a recent study of CD63-GFP mice, which estimated this to be ∼9-15% on the basis of SPIR-IFM imaging data. However, our paper is the first to use a genetic model to estimate the neuronal contribution to blood exosomes.

### Implications for EV-based diagnostics

Our detection of hepatocyte-derived and neuron-derived exosomes in the blood provides direct empirical support for the proposition that blood exosomes can report on the health and disease of both cell types. Furthermore, it indicates that further interrogation of cell type-specific exosomes may provide us with a more detailed understanding of the proteins and RNAs present in hepatocyte-derived and neuron-derived exosomes, and how their composition changes in different states of health (fed vs. fasted, sedentary vs exercise-trained, day vs night, etc.) and disease (diabetes, non-alcoholic fatty liver disease, alcoholic cirrhosis, traumatic brain injury, neurodegenerative diseases, etc.). This model might even be helpful in distinguishing between the opposing views of L1CAM as a marker of neuron-derived exosomes (85-88,95,96).

### Comparison to other transgenic models

Although the *exomap1* mouse was the first to be based on CD81 and the first to show the relative contributions of specific cell types to blood and CSF exosome population, it is not the first transgenic animal model to employ a highly-enriched exosome cargo protein, as previous studies have described animals that express CD9-GFP(97,98) or CD63-GFP(99–103). Of these, the TIGER mouse model is the most similar to the *exomap1* mouse, as it carries a Cre-activated, CAG-regulated transgene designed to express a fusion between full-length human CD9 and GFP (HsCD9-GFP(97)). Like the *exomap1* mouse, the TIGER mouse employs a highly enriched exosome cargo, CD9, as its exosome tracer protein, and moreover, uses the human form of CD9 so that marked exosome can be immunopurified on anti- HsCD9-specific monoclonal antibodies. Nevertheless, there are a few differences in transgene design between these two mouse models, as the TIGER mouse transgene (***i***) utilizes a silent lox-stop-lox cassette and therefore lacks an upstream reporter of pre-Cre transgene expression, (***ii***) was inserted randomly in the mouse genome instead of being inserted in a safe harbor locus, and (***iii***) is based on CD9, which is 3-fold less-enriched in exosomes than CD81(5).

In addition to the CD81-based *exomap1* and CD9-based TIGER mice, several groups have developed transgenic animals that express CD63 proteins tagged with a fluorescent protein (99–103). However, it is currently unclear whether these CD63-based models are the most appropriate tools for studying exosome biology *in vivo*, in part because CD63 is loaded into exosomes at ∼15-fold lower efficiency than CD81(5), in part because transgenic expression of CD63 is lethal(100), and in part because high-level expression of CD63 inhibits AP-2-mediated endocytosis, with complex effects on exosome content(44). Given that *in vivo* models for studying exosome biology should avoid perturbing cell physiology and exosome content, it is unclear whether CD63- based animal models are only mildly disruptive or induce major changes to cell physiology and exosome content.

### Limitations of the exomap1 model

We designed the *exomap1* mouse for expression in the broadest possible array of cell types by inserting it in the H11 safe harbor locus and using the CAG promoter. Furthermore, we demonstrated that the *exomap1* transgene is expressed in most cell types, including oocytes, cumulus cells, the corpus luteum, myeloid cells, skeletal muscle, muscle fiber peripheral cells, hepatocytes, neurons, and other CNS cell types. Nevertheless, our analysis of *exomap1* mice identified several instances in which we were unable to detect either MTS-tTomato or HsCD81mNG fluorescence. This issue highlights the value of including an upstream fluorescent marker in the *exomap1* transgene design, as it allows investigators to confirm expression of the *exomap1* transgene in specific cell types of interest, prior to embarking on Cre activation studies.

Another potential limitation of the *exomap1* mouse is suggested by the spurious release of reporter proteins by transgenic cells and animals(104–106). Specifically, it may be that HsCD81mNG-expressing cells might release some free mNG protein fragments into biofluids, and if so, this might confound the use of extracellular mNeonGreen fluorescence as a proxy measurement of HsCD81mNG release from the cell in exosomes. While this hypothetical problem might indeed complicate such an assay, we did not use this assay in our paper, and thus, this potential problem does not affect the utility of the *exomap1* model or the data presented in this paper. We nevertheless examined the mNeonGreen fluorescence of raw plasma fractions collected *exomap1*::*LysM*-Cre mice that had been separated by size exclusion chromatography. This revealed that most of the mNeonGreen fluorescence peaked in the exosome fraction (***fig. S4***). Thus, while some free mNeonGreen fluorescence may be released in soluble form, it does not impact the utility of the *exomap1* mouse as an *in vivo* model for studies of exosome biogenesis.

Finally, it should be noted that all Cre driver-based transgenic animal studies involve two general limitations. The first of these is that Cre drivers often display some degree of incomplete penetrance, which in the *exomap1* model would manifest as the failure to delete the MTS-tdTomato ORF and activated HsCD81mNG expression in cell type of interest. The second of these is that all Cre drivers are occasionally expressed in ‘off-target’ cells. These issues are unlikely to have had a significant impact on the data and conclusions of our study. In the case of animals infected with AAV8/*TBG*-Cre virus, this tool is known to drive Cre expression almost exclusively in liver hepatocytes due to the combined hepato-specific tropism of AAV8 and hepato-specific transcription from the TBG promoter(58,79–81,107). As for the *Camk2a*-Cre driver used in this study, it too has been subjected to extensive characterization that has demonstrated it drives Cre expression in broad array of neurons. As for the slight expression of this driver outside the CNS, this is due largely to its expression in peripheral nerves and to some spurious expression in the male germline(108). However, such spurious expression could only lead to an overestimation of neuronal contributions to blood EV populations, and thus would not affect our conclusion that neurons contribute <1% of the exosomes in blood and CSF. In short, while the limitations of Cre-mediated transgene expression are valid concerns, they are not a model-specific concern and do not impact the conclusions of our study.

### Data analysis and presentation

Statistical analysis involved calculation of averages and standard error of the mean, with pairwise differences evaluated for likelihood of null hypothesis using Student’s t-test, or ANOVA for experiments evaluating more than 2 sample sets. P-values for exosome diameters and marker protein abundance/EV from qSMLM data were calculated after logarithmic transformation to eliminate the bias of distribution skewness. Histograms and scatter plots were generated using Excel and Matlab R2022a. Images were imported into Adobe photoshop and figures were assembled in Adobe Illustrator. Image data was adjusted for brightness only.

## Supporting information

Supplemental material for Fordjour et al. Exomap1 mouse

## Data Availability

All data is contained within this paper.

## Resource availability

*Exomap1* mice will distributed through a commercial provider.

## Acknowledgments

We thank Drs. Michael Caterina and Shang Jui Tsai for assistance in maintaining the *exomap1* breeding colony at Johns Hopkins University, and Huizhen Wang for confocal imaging of the skeletal muscle tissues. Brain plate images in Figure 6 were generated with BioRender.

## Author contributions

Conceptualization: FKF and SJG

Data curation: SJG, TJ-T, LKC, SD, CDF, RLR, PCG

Formal analysis: SJG, TJ-T, LKC, SD, CDF, RLR, PCG

Funding acquisition: SJG, TJ-T, LKC, SD, CDF, RLR, PCG

Investigation: FKF, SJG, SA, XH, EC, VL, MN, AS, FD, NC-A, HW, KS, MB, TAP, NKV, PCG, RLR, CDF, SD, LKC, TJ-T

Methodology: SJG, TJ-T, LKC, SD, CDF, RLR, PCG

Project administration: SJG, TJ-T, LKC, SD, CDF, RLR, PCG

Resources: SJG, TJ-T, LKC, SD, CDF, RLR, PCG

Supervision: SJG, TJ-T, LKC, SD, CDF, RLR, PCG

Validation: SJG, TJ-T, LKC, SD, CDF, RLR, PCG

Visualization: FKF, SJG, SA, XH, EC, VL, MN, AS, FD, NC-A, HW, KS, MB, TAP, NKV, PCG, RLR, CDF, SD, LKC, TJ-T

Writing – original draft: SJG

Writing – reviewing and editing: FKF, SJG, SA, XH, EC, VL, MN, AS, FD, NC-A, HW, KS, MB, TAP, NKV, PCG, RLR, CDF, SD, LKC, TJ-T

## Funding

This work was supported by the National Institutes of Health to S.J.G. (U19 CA179563 and UG3 CA241687), L.K.C. and P.G.C. (R21 AG066488), S.D. (R35 HL150807), C.D.F (R01 DA051831 and DP1 DA039658), T.J.-T. (UG3/UH3 TR002878), and R.L.R (UG3/UH3 CA241703). R.L.R. was also supported by the Department of Veterans Affairs (I01BX003928 and IK6BX005692). The content of this paper is solely the responsibility of the authors and does not necessarily represent the official views of the National Institutes of Health.

## Conflict of interest

F.K.F. and S.J.G. are potential beneficiaries of licensing and other fees related to use of *exomap1* mice, which are owned by Johns Hopkins University. The remaining authors declare that they have no conflicts of interest with the contents of this article.

