## Supplemental material for Fordjour et al. Exomap1 mouse for "Exomap1 mouse: a transgenic model for *in vivo* studies of exosome biology"

**Supporting information**

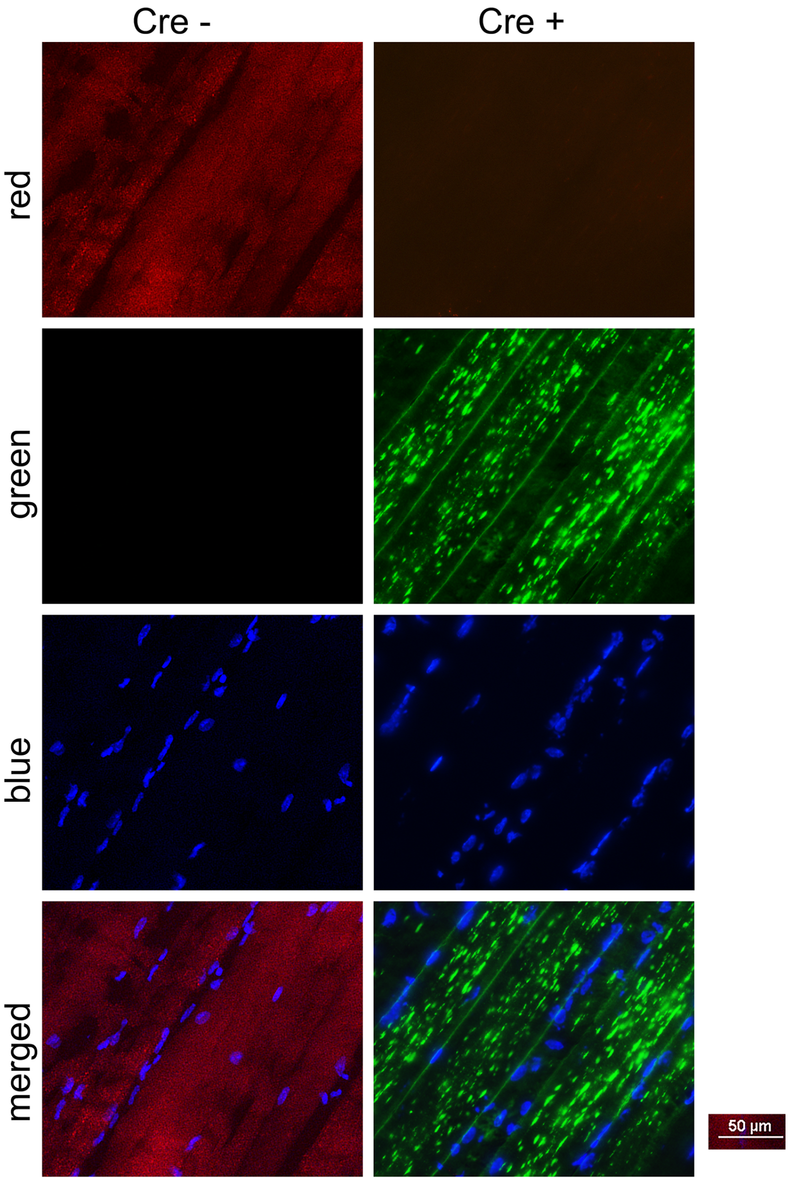


*Figure S1. Cre-induced expression of HsCD81mNG in skeletal muscle.* Fluorescence micrographs of DAPI-stained quadriceps muscle sagittal sections showing the Cre-mediated switch of *exomap1* transgene products from MTS-tdTomato to HsCD81mNG between *exomap1*$^{+/}$ mice and tamoxifen-treated *exomap1*::*HSA*-Cre$^{ERT2}$ mice.

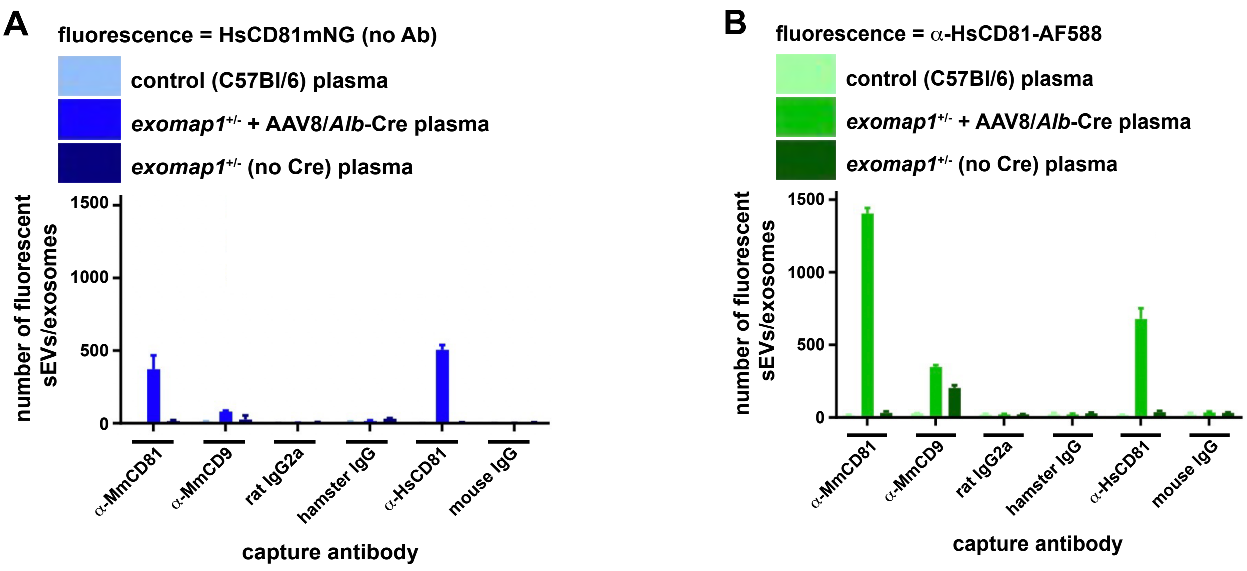


*Figure S2. SPIR-IFM analysis of serum exosomes from control, exomap1*$^{+/}$*, and exomap1::AAV8/TBG-Cre mice.* Histograms of SPIR-IFM imaging data show the numbers of fluorescently-labeled exosomes captured from serum (control animals, animals expressing HsCD81mNG in the liver, or *exomap1* animals not exposed to Cre), bound to different capture antibodies specific for MmCD81, MmCD9, HsCD81, and controls. (**A**) Bar graph of HsCD81mNG fluorescence on plasma exosomes captured on different antibodies. (**B**) Bar graphs of Alexa Fluor 555 anti-CD81 staining on exosomes captured on different antibodies.

In brief, plasma exosomes were immunopurified on SPIR imaging chips functionalized with antibodies specific for MmCD81, MmCD9, and HsCD81, as well with three control antibodies (IgG from rat, hamster, and mouse). Once captured, the immobilized sEVs were incubated with an Alexa Fluor 588 (AF588)-labeled antibody specific for HsCD81, washed, dried, and examined by SPIR-IFM imaging. The resulting images showed that green HsCD81mNG fluorescence was detected only in serum exosomes from *exomap1*$^{+/}$ mice infected with the liver-specific, Cre-expressing virus (***fig. S2A***). Immunostaining for HsCD81mNG supported this conclusion, as it revealed that hundreds of the exosomes captured on MmCD81 antibodies also stained positive with the HsCD81 antibody (***fig. S2B***).

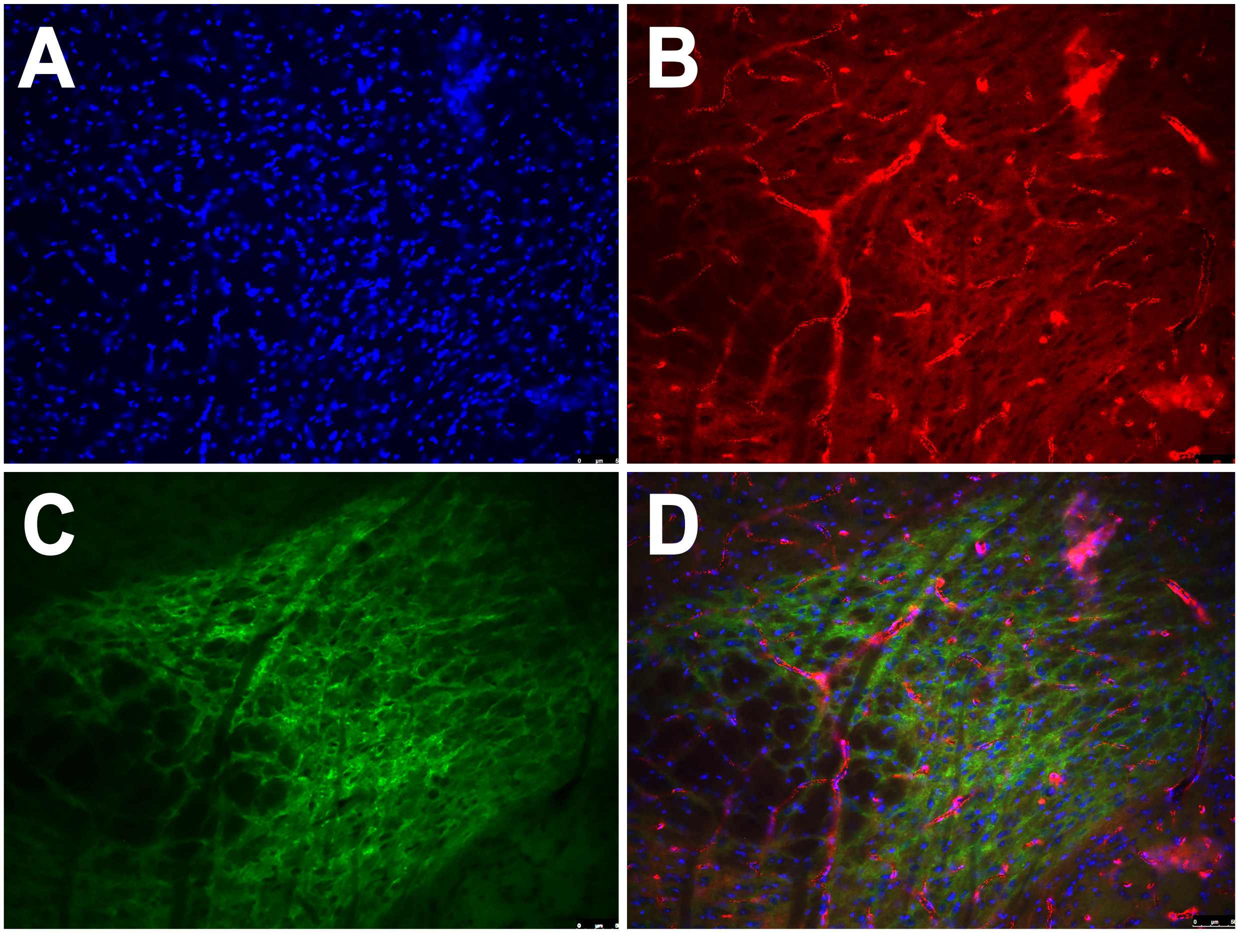


*Figure S3. Dat-Cre activates expression of HsCD81mNG in midbrain*. Fluorescence micrographs of DAPI-stained brain section showing fluorescence of (A) DAPI, (B) MTS-tdTomato, (C) HsCD81mNG, and (D) a merge of all three. Bar, 50 um.

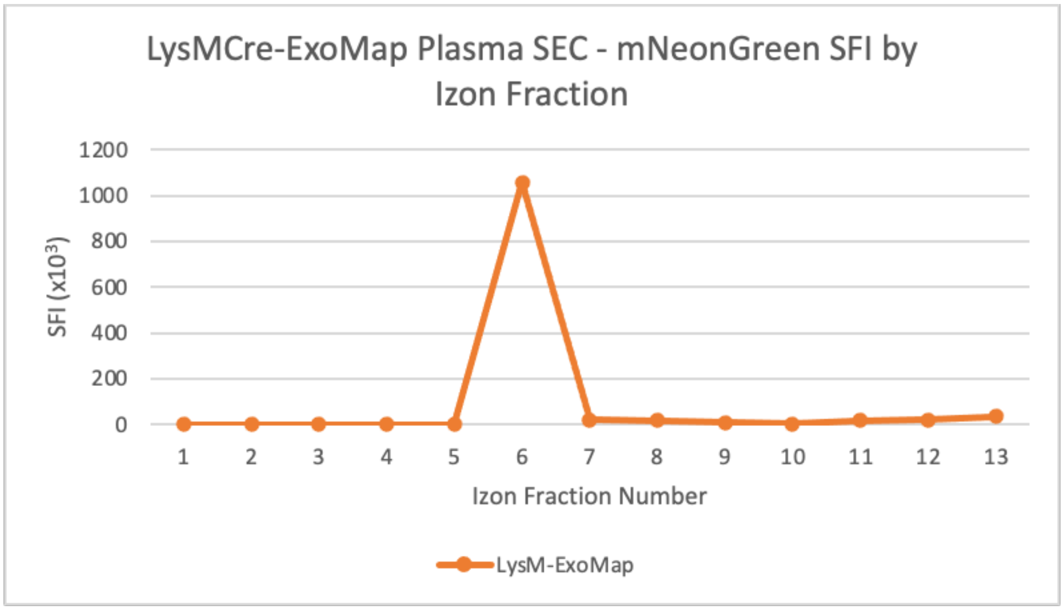


*Figure S4. Plasma mNeonGreen fluorescence* exomap1*::*LysM-*Cre mice elutes in the peak exosome fraction*. Bar graph of specific mNeonGreen fluorescence intensity (SFI; relative brightness (arbitrary units)/protein concentration) in fractions of plasma samples. Raw plasma from *exomap1*::*LysM*-Cre mice was separated the plasma by size exclusion chromatography (SEC) using an Izon qNano column (Izon Sciences). Each fraction was assayed each fraction for mNeonGreen fluorescence and protein concentration. Void volume (fractions 1-5) is followed by EVs (fraction 6) and free proteins (fractions 8-13).

**Table S1. Summary of p-values for the average diameter and number of detected TSPAN molecules per vesicle**.

| ***Exomap1***$^{+/:}$**::AAV8/*TBG*-Cre** | **p-value TSPAN/vesicle** | **p-value vesicle** $D_{Av}$ |
|---|---|---|
| MmCD81 capture, stain with MmTSPAN Abs vs HsCD81 Ab | <0.001 | <0.001 |
| HsCD81 capture, stain with MmTSPAN Abs vs HsCD81 Ab | <0.001 | 0.2 |
| HsCD81 capture, HsCD81 stain vs MmCD81 capture, MmTSPAN stain | <0.001 | 1.0 |

| ***Exomap1***$^{+/:}$**::Camk2a-Cre** | **p-value TSPAN/vesicle** | **p-value vesicle** $D_{Av}$ |
|---|---|---|
| HsCD81 vs MmCD81 capture, plasma | <0.001 | <0.001 |
| HsCD81 vs MmCD81 capture, CSF | 0.9 | 0.2 |
